# Forecasting Extinction Risk using Thermal Performance Curves and Population Dynamic Modeling

**DOI:** 10.1101/2025.04.27.650737

**Authors:** David A. Vasseur, Carling Bieg, Misha Kummel, Alison J. Robey

## Abstract

Thermal Performance Curves (TPCs) have become a popular tool for assessing the risks imposed by climate warming and variability on ectotherms. These assessments typically measure the match between an organism or population’s TPC and the distribution of its current or future thermal environment as a proxy for extinction risk. However, extinction can occur even when the average thermal environment appears closely matched to a population’s TPC because population dynamics can be very sensitive to thermal stress. Here, we develop a new metric for assessing extinction risk using a stochastic model of logistic growth as a foundation. We show that boundaries delimiting persistence and extinction regions of parameter space can be derived for the simple case where the intrinsic (Malthusian) growth rate *r* varies stochastically and that these boundaries continue to make reliable predictions when temperature *T* varies stochastically and the Malthusian growth rate is given by a thermal performance curve *r* (*T*). We accomplish this by combining theory with stochastic simulations of population dynamics and a laboratory experiment where populations of the single-celled protist *Paramecium caudatum* were cultured across different temporal means and variances of temperature. The measure of risk that we develop and validate is straightforward and easily applicable to any population for which the thermal performance of Malthusian fitness is known, allowing more rigorous identification of the risks imposed by warmer and more variable temperatures across the globe.

---

Thermal Performance Curves (TPCs) have become a common tool for assessing the suitability of environmental conditions and the impact of warming on ectotherms (*1–5*). Typically, these curves are left-skewed unimodal functions of temperature that decline to zero at thermal extremes when performance metrics are biological rates such as locomotor speed, foraging or ingestion rate, maturation rate, or lifetime reproductive output (*6*). However, when combined into aggregate processes such as the population growth rate (Malthusian fitness, *r*), TPCs descend below zero at the thermal extrema, reflecting a deficit in births relative to deaths, and leading to population decline (*3, 7, 8*). Many authors have used TPCs to forecast the inherent risk of climate change for populations or species by comparing their TPC to the historical, current, and/or future thermal environment. Some studies have focused on the number or frequency of heat events that exceed a critical temperature (*1, 5*), while others have calculated the change in long-term average fitness of a population (*2, 3*). Unfortunately, neither of these measures have a clear, mechanistic relationship with the most important measure of risk: the probability of extinction. Although a population enduring temperatures that lead to a negative growth rate over the long term (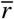 < 0) will certainly be driven to extinction, even populations with a positive mean growth rate (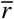 > 0) can have a high risk of extinction when exposed to frequent or prolonged periods of stress (defined as those during which *r* < 0). To properly address extinction risk, we must recognize that extinction is a characteristic outcome of population dynamics and thus any attempt to make inferences about the connection between environmental variation and extinction risk will benefit from a population dynamic framework.

The impact of environmental stochasticity on the risk of population extinction has long been a topic of investigation, with early work by Levins (*9*) and May (*10*) focused on how variation in the parameters *r* and *K* of the logistic model impacted population dynamics and the risk of extinction. Similar work occurred even earlier in the field of population genetics, albeit with a focus on the roles of selection and drift rather than persistence (*11*). Lande (*12*) extended this work to consider the impact of demographic stochasticity and random catastrophes, and more recent work has explored the role of autocorrelation (*13–15*). Most forms of stochasticity lead to greater variability in population dynamics, thereby increasing the risk of extinction, but environmental variation also tends to reduce average population sizes due to non-linearities in the functions governing population growth (*16*). For example, when the carrying capacity (*K*) of a Logistic model varies sinusoidally, population size approaches the smaller harmonic mean, rather than the arithmetic mean of *K* (*10, 17*).

Temperature variation can interact with the non-linearity of TPCs to generate a surprisingly broad range of outcomes. Siddiqui (*18*) demonstrated that aphid populations alternating between two temperatures grew more rapidly than populations grown at the mean (constant) temperature. This difference arises due to non-linear averaging (Jensen’s inequality) and has been central to the discussion on temperature variability. In general, temperature variation can substantially depress the average population growth rate at warmer temperatures where the TPC is a concave down function (e.g. (*3, 19*)) which has led to the observation that the modal temperature of TPCs should be greater than the environmental average to offset the negative effect of variability (*20*). Nonetheless, the effect of temperature variation on population performance and extinction risk has not been quantified in a generalizable way.

Duffy et al. (*21*) recently used a model of temperature-dependent population dynamics to measure the risk of extinction using a well-studied set of TPCs from 38 terrestrial invertebrate ectotherms that were first described by Frazier et al. (*22*) and subsequently used in many studies (*2, 3, 23*). Although they found that projections of nearly all populations were more prone to extinctions in future climate scenarios, the mean, variability, and autocorrelation of population dynamics (independently and in concert) were not useful proxies for extinction risk. This highlights the challenge of identifying the impacts of changing climate based solely on trends in population abundance and other characteristics of their dynamics.

Here, we provide an important advance in estimating the risk of warming and temperature variation. We couple an analytically derived measure of extinction risk from a simple stochastic model of logistic population growth with recent work that specifically considers the integration of temperature into this model (*8, 21, 24*). We demonstrate that this measure of extinction risk is a robust predictor of the risk imposed by temperature variation even though the non-linearity of thermal performance curves violates a key assumption (normality) of our analytical metric. We validate our findings in a microcosm experiment where *Paramecium caudatum* was grown under different thermally variable conditions. Importantly, this metric of extinction risk can be derived without any additional information beyond those already commonly used to estimate risk: the TPC and the distribution of temperature experienced by the population.

## Extinction in simple stochastic environments

A useful starting point for introducing environmental variation into population dynamics is the *r − α* model,

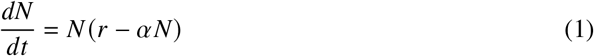

an analogue of the logistic model where the intrinsic growth rate, *r*, is separated from the density-dependent term, *αN*, in the per-capita growth rate [see (*25*)]. Notably, the effects of temperature are similarly separated across the two terms; *r* has been shown to follow a TPC while *α* is often assumed to be independent (see below).

The non-linearity of TPCs makes a temperature-dependent version of Equation 1 difficult to analyze so we begin by assuming a more generic form of environmental variation which leads to temporal variation in *r* that is normally distributed (IID - independently and identically distributed), with mean 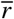 and variance 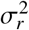. This model can be written as the stochastic differential equation:

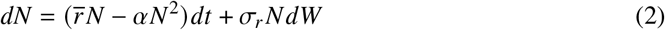

where *W*, the Wiener process, is used to generate stochastic increments in continuous time (see Supplementary Text). For this model, the stationary probability density of population size *p* (*N, t*^*^) can be analytically solved by diffusion approximation (*9, 10, 26*). *p* (*N*) has the form of a Gamma distribution and undergoes two critical changes as the ratio 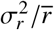 increases; these have important impacts on the probability of extinction. For values of 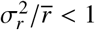, the distribution *p* (*N*) is unimodal and bounded away from zero. Here, extinction is very unlikely because periods of stress (when *r* < 0) are relatively uncommon compared to periods conducive to growth (though note that density dependence may lead to population declines even when *r* > 0, if *N* is sufficiently large). But when 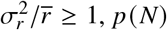 is a monotonically decreasing function of *N* with the greatest probability density at *N* = 0. May (*10*) suggested that this threshold could be taken as the boundary separating likely persistence from likely extinction and herein we term this the ‘weak persistence boundary’. Beyond this boundary, periods of stress are frequent enough that the population can become trapped at low densities, increasing the risk of extinction. A second ‘strong persistence boundary’ exists at the critical point 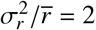, where the entire probability distribution *p* (*N*) collapses into a single point at *N* = 0. Above this strong boundary, populations can persist only in short-term transient states, if initialized at high density. Here, the time to extinction may vary but extinction itself is a certain outcome.

In differential equation models like Equation 1 and 2, extinctions are never truly realized because population densities only approach extinction, *N* = 0, in the limit of infinite time. However, by setting an arbitrary extinction threshold *N*_*e*_ = 1, we can calculate the mean time required for the population to reach extinction (*t*_*e*_) given an initial abundance *N*_0_. At the weak boundary, *t*_*e*_ *≈ α*^*−*1^ for reasonably large *N*_0_; around the weak boundary, *t*_*e*_ changes non-linearly as a function of 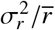, leading to rapid extinction above the boundary and prolonged persistence below it (see Supplementary Text).

Fig. 1 shows the location of the weak and strong boundaries overlaid on results from numerical simulations of Equation 2 after *t* = *α*^*−*1^ time steps have elapsed. As expected, approximately half of the simulated populations are extinct at the weak boundary, while nearly all of the populations are extinct at the strong boundary and beyond. Sample probability density functions from the three regions delimited by the weak and strong boundaries show the expected transition from unimodal, to monotonically decreasing, to non-existent (Fig. 1b-d). Here, it is clear that the weak persistence boundary is a robust predictor of extinction risk.

**Figure 1:**
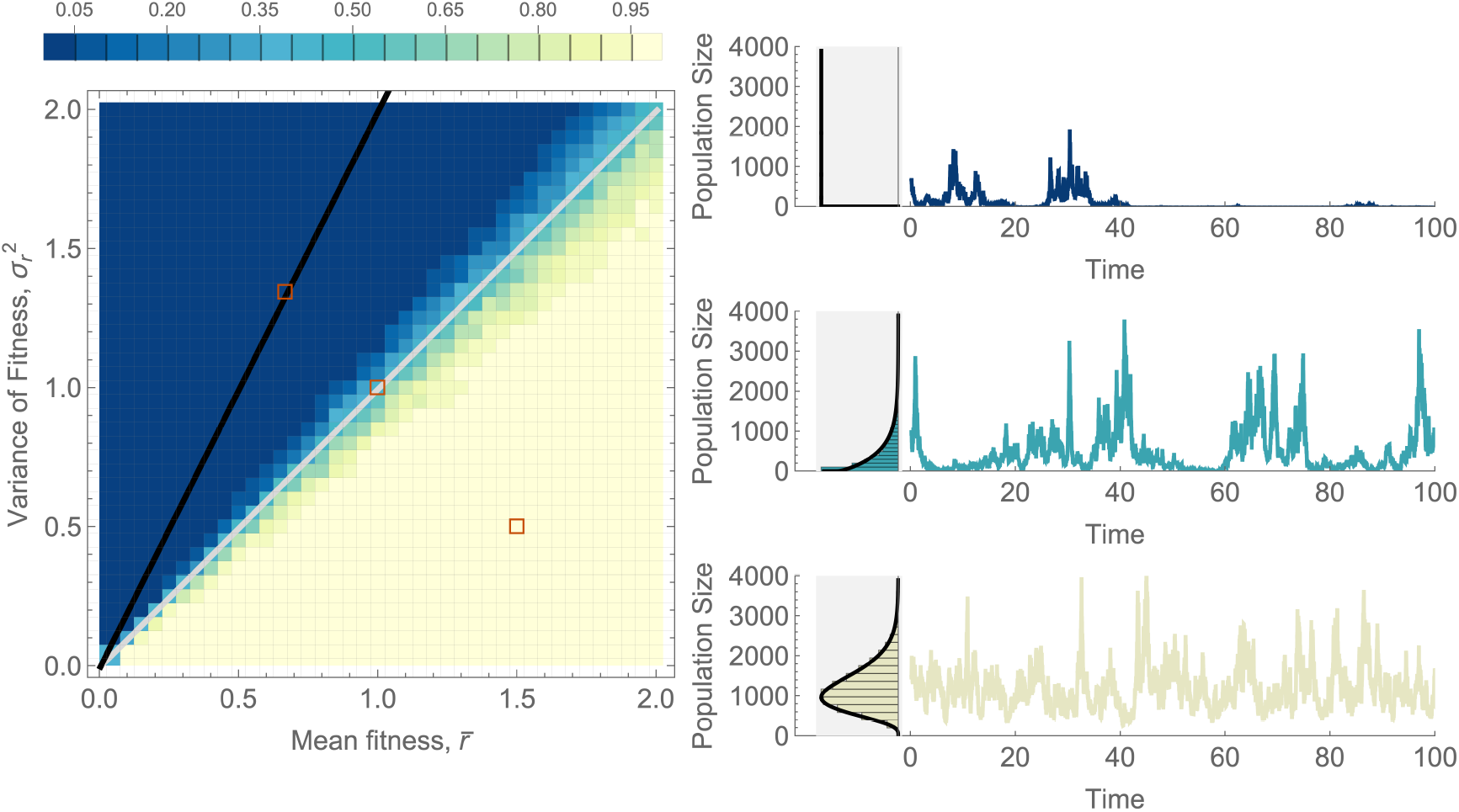
The fraction of replicate simulated populations persisting after *α*^*−*1^ = 1000 time-steps for the stochastic *r − α* model. The strong persistence boundary, 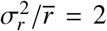, is shown in black and the weak persistence boundary, 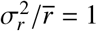, is shown in gray. Sample population dynamics and their sample (bars) and predicted (black lines) probability distributions are shown for three reference points (red boxes) in the parameter space. For each parameter pair, we initialized 100 populations at 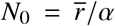 and numerically integrated the stochastic differential equation using Milstein’s method (*37*) (Supplementary Text). We assumed that a population was extinct if it dropped below *N*_*e*_ = 1. We use *α* = 0.001 and show in the Supplementary Text how the choice of *α* affects our result. A time-elapsed animation of this figure is available in Supplementary Movie S2.

## Extinction due to temperature stochasticity

Numerous studies have considered the role of temperature variation in population dynamic models (*8, 21, 24, 27*) and there is broad agreement that in ectotherms, *r* (*T*) follows a unimodal, typically left-skewed TPC (*6, 28*). Less is known about how temperature modifies the strength of density-dependence (*8*), but recent evidence suggests a stronger density dependence at higher temperatures due to the larger per-capita resource requirements needed to offset higher rates of respiration (*24*,*29*). For simplicity, here we assume that density dependence does not change with temperature; this leads to an equilibrium abundance that also tracks a thermal performance curve given by *r* (*T*)/*α*. Given that we have outlined expectations for how variation in *r* influences extinction risk (see above), it is tantalizing to hope that those expectations will hold when temperature drives variation in *r* via the thermal performance curve. Unfortunately, even a simple normal distribution of temperature variation introduces skewness, kurtosis, and bias to higher moments of the distribution of *r* due to the non-linearity of TPCs (Fig. 2), thereby violating the assumptions used to construct the weak and strong boundaries above and potentially reducing the predictive utility. In the sections that follow, we show that these boundaries continue to be surprisingly robust predictors of extinction risk for populations experiencing temperature variation.

**Figure 2:**
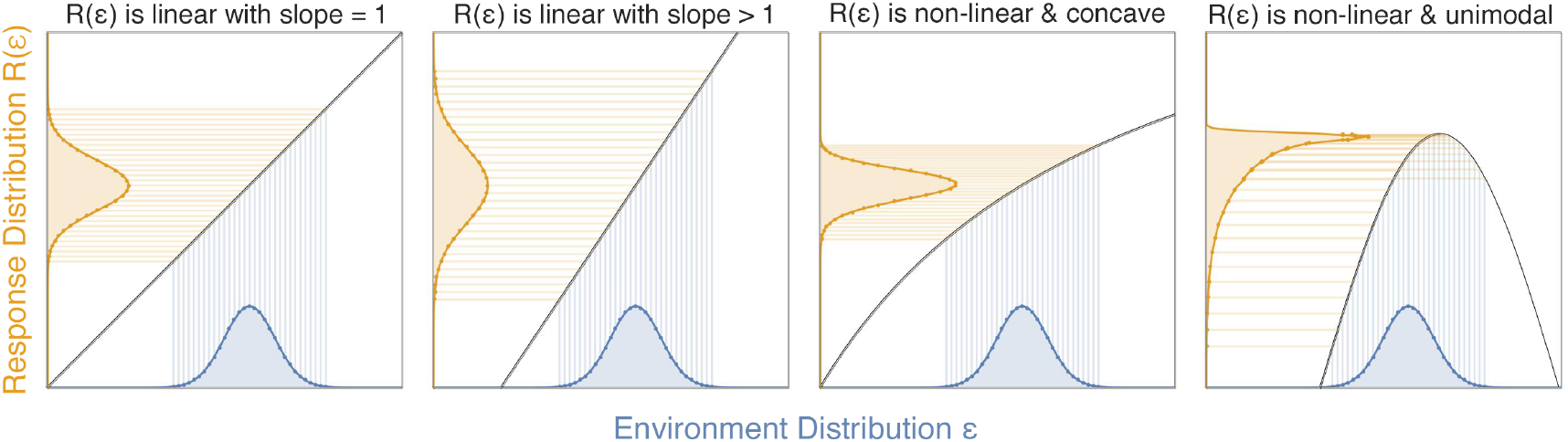
The transformation of a normally-distributed environmental variable through four different response functions. When the function *R*(*ϵ*) is linear (panels a and b) the mean and variance can be altered (the latter only when |*dR*/*dϵ* | ≠ 1), but higher moments are unaffected. When the function *R*(*ϵ*) is non-linear (panels c and d), the mean,variance, and higher moments (skewness, kurtosis, etc.) are altered. Thermal performance curves are most similar to the rightmost panel, and lead to highly skewed and leptokurtic distributions of fitness when temperature is normally distributed.

Assuming that temperature is a normally distributed random variable 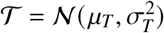, we find the locations of the weak and strong boundaries by taking expectations:

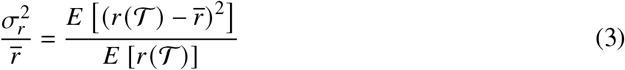

In practice, this requires numerical approximation for most, if not all, functional representations of the TPC. In Fig. 3, we compare the weak and strong boundaries to simulation results from a hybrid SDE (*r* (*T*) *− α*) model, where *r* (*T*) is given by the TPC of the unicellular ciliated protist *Paramecium caudatum* (see Supplementary Text). When translated onto axes describing the distribution of temperature, the weak and strong boundaries enclose a region of likely persistence that is widest in the absence of thermal variation and narrows asymmetrically as thermal variation increases due to the strong negative effects of experiencing more temperatures above *T*_max_. Despite the fact that the weak and strong boundaries assume a normal distribution of *r*, they show a remarkably good coherence to simulation results, even at large values of *σ*_*T*_ where the distribution of *r* shows strong skewness and kurtosis (Supplementary Text). Examination of the population dynamics shows that they retain much of their characteristic behavior, with the exception that the dynamics between the weak and strong boundaries are bounded away from zero, explaining the lower than expected extinction risk in this range (Fig. 3d). Compressing the simulation results from all different combinations of *μ*_*T*_ and *σ*_*T*_ onto a single axis defined by 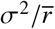 in Equation 3 shows that this ratio is a strong predictor of extinction risk, and that the weak boundary where 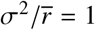 indicates the point where extinctions begin to occur on the 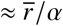 timescale.

**Figure 3:**
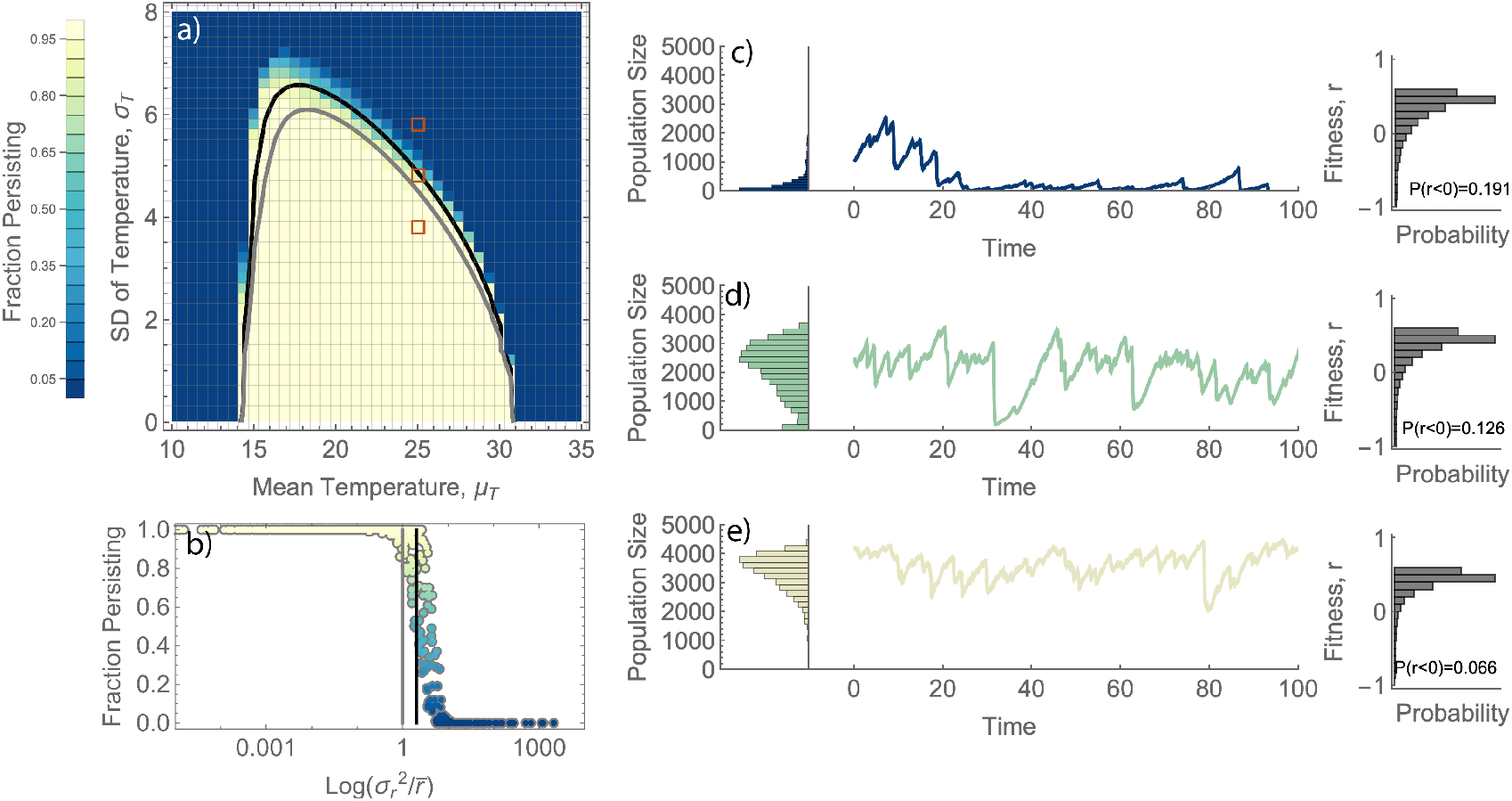
The fraction of replicate populations persisting after 10000 time-steps for the *r* (*T*)*−α* model. (**a**) Simulated results when temperature variation is normally distributed 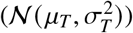 The **strong persistence boundary**, 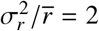, is shown in black and the **weak persistence boundary**, 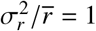, is shown in gray. (**b**) the data from (**a**) are aggregated onto a single axis, 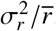, showing that an extinction cascade begins at the weak boundary regardless of the specific attributes of the environment. (**c-e**) Population dynamics and their sample probability distributions are shown for three reference points in the parameter space (red boxes in **a**) along with the probability distribution of fitness, *r*. For each parameter set we initialized 100 populations at 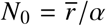 and numerically integrated a hybrid SDE with *α* = 0.0001. We assumed that a population was extinct if *N* dropped below 1 individual. A time-elapsed animation of this figure is available in Supplementary Movie S2.

To further test the predictive utility of 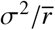, we grew 12 replicate cultures of *P. caudatum* for 8 weeks in thermal environments that changed every 12 hours across 20 different combinations of thermal means and variances. The fraction of cultures that persisted in each treatment combination exhibited a strong concordance with the location of the analytical boundaries, which were calculated using only the independently measured TPC (Fig. 4).

**Figure 4:**
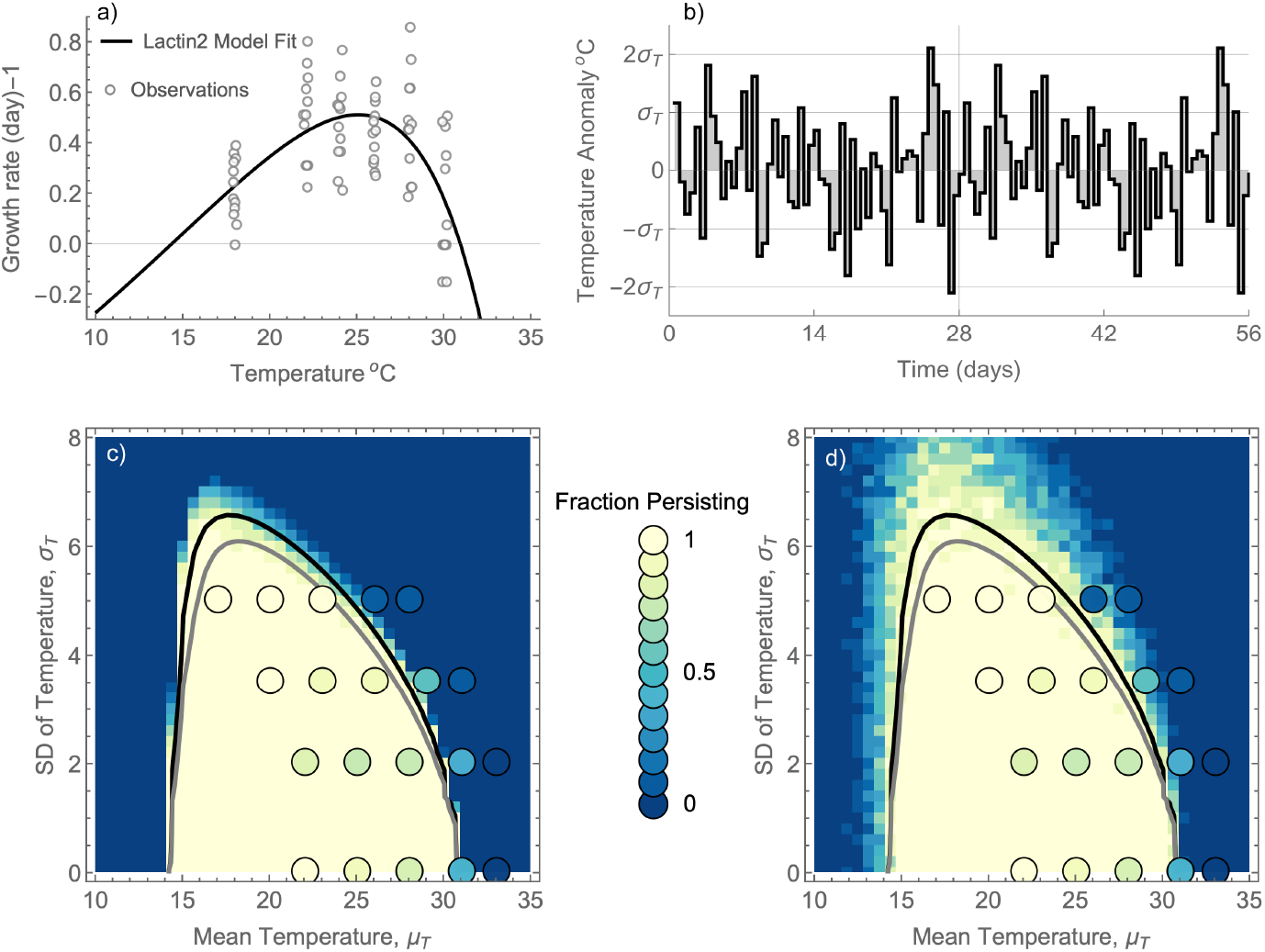
Extinctions in *P.caudatum* cultures are accurately forecast by the persistence boundaries calculated using the thermal performance curve *r* (*T*). (**a**) The TPC of *r* fit to experimental observations using the lactin2 model (*38*). (**b**) shows the temperature anomaly that was used to generate (via scaling and translation) each of the 20 temperature series used in the extinction experiment (note that the sequence repeated after 28 days (**c, d**) show the fraction of experimental populations persisting at the end of the experiment (embedded circles) superimposed on simulation results from the SDE model (c) and from a stochastic simulation model which followed the exact temperature sequences from our experiment and included the effects of demographic stochasticity and decaying culture conditions (d). Above (below) the strong extinction boundary 71/88 (13/137) replicates went extinct (15 replicates were discarded due to low/no abundance at the beginning of the experiment). Using a permutation test we found that extinctions were significantly more likely to occur in populations that were positioned on or above the strong persistence boundary (*p* < 10^*−*6^).

In our experimental cultures, we estimate the strength of density dependence to be *α ≈* 0.0001 by dividing the observed intrinsic growth rate *r* (*T*) *≈* 0.5 by the observed equilibrium population size *≈* 5000 cells at *T* = 25°C (unpublished). This results in a prediction of the mean time to extinction on the order of *α*^*−*1^ = 10, 000 days. Our experiment was much shorter than this, but several potential factors explain why the model still predicted extinction risk well. Firstly, the mean time to extinction predicted by the model changes rapidly in the neighborhood of the weak boundary; moving only slightly beyond the boundary can result in mean times to extinction that are 1-2 orders of magnitude smaller. Our model also omits several important processes that are fundamental drivers of extinction in our experiments: (i) The demographic stochasticity present in our experimental populations is not incorporated in our SDE model or persistence boundaries. Demographic stochasticity should decrease the mean time to extinction in all environments, thereby increasing the number of extinctions occurring in our experiment. (ii) Temporal autocorrelation is absent from the theory used to generate persistence boundaries, though it permeates both our simulation and experimental results in Figures 3 and 4. Temporal autocorrelation increases extinction risk by lengthening periods of population decline relative to uncorrelated variation. In our model simulations, autocorrelation was insignificant beyond the daily scale; in our experiment, temperature was constant within 12 h intervals, but uncorrelated across steps. (iii) Conditions in our batch-culture experiment decay over time, leading to lower population sizes and higher rates of extinction over time. We constructed a stochastic simulation model that mimicked the conditions in our experiment, including demographic stochasticity, abrupt temperature changes every 12 hours, and a doubling of *α* over the course of the 56 day experiment. Figure 4d shows that the addition of these factors understandably make the simulation results more noisy, but aligns the simulation and experimental results to a similar time-scale.

## Extensions and Limitations of our approach

For any population for which *r* (*T*) has been estimated, we can easily calculate the position of the weak and strong boundaries for given values of *μ*_*T*_ and 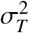 (e.g., Fig. 3c). This boundary encloses an environmental parameter space where persistence is likely, much like the Hutchinsonian niche. Holt (*30*) defines the Hutchinsonian niche as the environmental space where *r* > 0; we extend this concept to show that the inclusion of temporal variability further limits the scope of that niche, since the persistence boundary 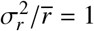 will always fall within the environmental space enclosed by *r* > 0. Measuring the proximity of a population to the persistence boundary (e.g. as 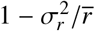) should provide a more conservative and robust measure of the relative risk imposed by a particular thermal environment than prior techniques focused on the (mis)match between the TPC and the thermal environment. One such measure of (mis)match that has been commonly employed is simply 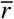, which has shown a decrease (increased risk) for tropical species, but an increase for temperate species (*2*). Our work demonstrates that extinction risk may increase even when 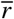 rises with climate change if altered climates also increase 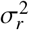. More work is needed to understand the connections between TPC shape, 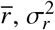, and the characteristics of natural temperature variation.

Forecasting extinction risk with TPCs is complicated by a number of factors (see (*31*)), including a complex relationship between operative body temperatures and ambient conditions (due to behavioral thermoregulation, microclimate variation, and other factors). Furthermore, thermal performance can change during acute and chronic exposure due to acclimation (*32*) and organisms can express vastly different responses to temperature during different ontogenetic stages or during periods of dormancy (*33*). The demographic buffering hypothesis states that for populations with age or stage structure, selection should reduce variation in the vital rates with the greatest impact on fitness (*34, 35*). Our work reiterates this idea, demonstrating that the most robust populations will be those that maintain high average fitness while buffering temporal environmental variation to ensure a low variance in fitness. In practice, this is difficult to achieve because thermal performance curves may be constrained by a generalist-specialist type tradeoff (*6, 36*), implying that broader TPCs capable of reducing variation in fitness would yield lower average fitness. In fact, given the canonical unimodal shape of TPCs, there are few situations where changing environmental conditions could lead to a simultaneous reduction in variance and increase in mean fitness. The extent to which TPCs are shaped by evolution or individuals are governed by thermoregulatory behavior to minimize the ratio 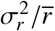 is worthy of further study.

We have shown that a simple integration of thermal performance into a population dynamic model leads to outcomes (persistence and extinction) that are rigorously predicted by analytical solutions obtained under simplified assumptions and confirmed using microcosm experiments. The ratio given by 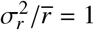 can be determined in a natural or simulated environment for any population whose thermal performance curve, *r* (*T*), is known, and its proximity to the persistence boundary can be easily quantified as a metric of extinction risk. Although more work is needed to probe how the metric performs under varied assumptions about the structure of population dynamics and thermal environments, this work provides an important advance in our ability to broadly assess the risk imposed by warmer and more variable climates.

## Acknowledgments

William Brown and Kai Padilla-Smith sampled and maintained *P. caudatum* communities and W. B. experimentally constructed its thermal performance curve.

## Author contributions

DV constructed and analyzed the stochastic differential equation models, designed the experiment, and led in writing the manuscript. AR, CB, and MTK contributed to various aspects of the simulation models, oversaw the experimental work, and contributed to writing the manuscript.

## Data and materials availability

Data and code are available upon request

## Supplementary materials

Materials and Methods

Supplementary Text Figs. S1 to S5

Table S1

References *(7-44)*

Movie S1 and S2

## Supplementary Materials

### Materials and Methods Summary

#### Simulations

We simulated the *r −α* SDE model (Equation 2) using the Milstein method (*37*) assuming a time-step of Δ*t* = 0.01. For all simulations, we set the initial population size as 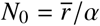 and assumed that an extinction occurred if *N* < 1 at any point during the simulation. To embed temperature variation into the *r − α* framework, we assumed that *r* varied as a function of temperature according to a left-skewed thermal performance curve given by the “lactin2” model (*38*) fit to our experimental system (see below) and we simulated a hybrid SDE where temperature varied according to an Ornstein-Uhlenbeck (OU) process:

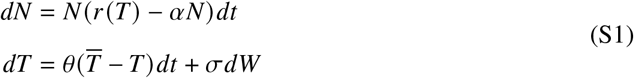

Since the OU model generates autocorrelated dynamics, we set the reversion speed to a large value (*θ* = 5) to ensure that the timescale of autocorrelation was much shorter than the generation times of our population. This ensured the closest possible match between the assumptions of our analytical boundaries and temperature-dependent *r* (*T*) *− α* model. Finally, we constructed a stochastic simulation model matching the specific details of our experiment and including demographic stochasticity. Briefly, we constructed temperature-dependent rates of birth and death for *P. caudatum* that resulted in the same net growth function *r* (*T*) we measured, then simulated the model using the Gillespie Stochastic Simulation Algorithm (*39*) following the same environmental conditions encountered in our experiment. Details on model simulations are given in Appendices C-E.

#### Experiment

We measured the thermal performance curve of *Paramecium caudatum* (Carolina Biological Supply) following the method outlined in (*27*) by seeding 12 replicate cultures with a known number of cells (between 4 and 6) into 1.5 mL of culture medium on a 24-well plate at 6 different temperatures. We incubated cultures for 48 hours, counted the number of cells, and calculated *r* for each replicate population. We then fit the “lactin2” model TPC to the data in *R* using the *rTPC* package (*38*). This fitted curve is shown in Figure 4 and is the basis for all our model simulations.

Using the weak and strong boundaries predicted from the TPC using Equation 3, we generated 20 experimental temperature sequences across a gradient of thermal means and variances and grew 12 replicate populations of *P. caudatum* in 14 mL culture tubes in each environment for 56 days, with temperatures changing randomly according to the generated sequences every 12 hours and repeating after the first 28 days. At the beginning, 28 day mark, and end of the experiment, we assayed for persistence/extinction. During first 28 day period, our heating apparatus was found to be were under-performing by approximately 2.5°C. We made adjustments on day 28 so that the second 28-day period reached the expected mean temperatures. Further experimental details are given in the Supplementary Text.

## Supplementary Text

## Appendix A The effect of stochasticity in *r* in the *r − α* logistic model

Beginning with the *r − α* model of population dynamics:

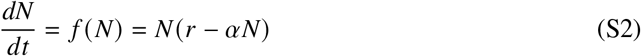

we are interested in knowing the effect of temporal variation in *r* on the dynamics and likelihood of extinction of *N*. This model has been studied in this same context by May (*10*) and by Pasquali (*26*). In the absence of variation, this model has equilibrium points at *N*^*^ = 0 and *N*^*^ = *r*/*α*. The latter is globally stable when *r* > 0 and the former whenever *r* < 0, with *r* = 0 representing a degenerate point where all values of *N* have neutral stability.

To incorporate uncertainty in *r* we introduce the auxiliary process *y*:

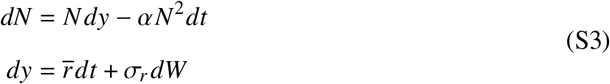

where 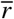 represents the mean value of *r, σ*_*r*_ is the standard deviation of *r*, and *W* is the Wiener process. Recombining the equations in (S3) gives the following stochastic differential equation:

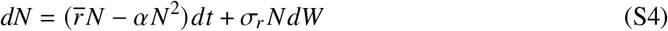

which is an Itô process, separated into drift and diffusion terms, respectively. From Equation S4, the probability density of population size *N, p* (*N, t*), can be estimated using the Fokker-Planck (or forward-Kolmogorov) equation:

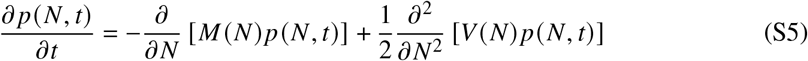

where *M* (*N*) and *V* (*N*) represent the mean and variance of *dN*/*dt*. The stationary probability density function, if it exists, is found by setting Equation S5 equal to zero and solving for *p* (*N, t*^*^). Wright (*40*) showed that the solution of *p* (*N, t*^*^) can be written as (see also the work of Levins (*9*) and May (*10*)):

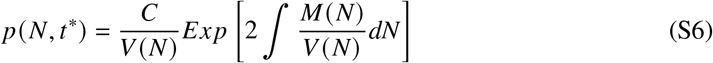

where *C* is a normalization constant given by:

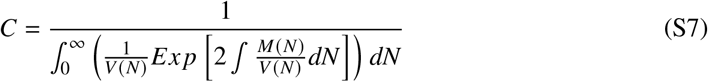

Taking expectations from Equation S2, we find:

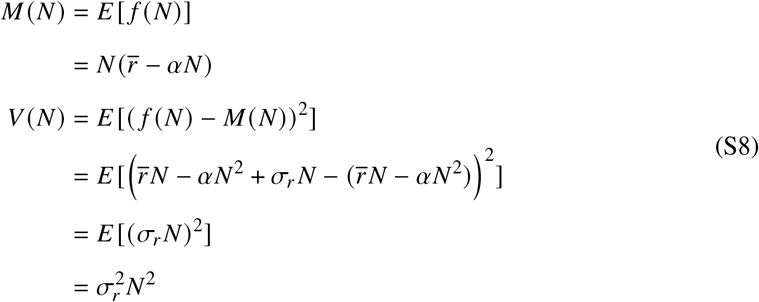

and solving the integrals in Equation S7 and setting 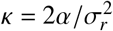 and 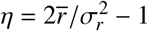 yields:

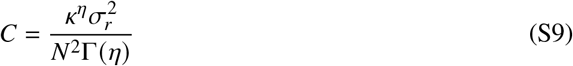

where Γ(·) is the Gamma Function. Substituting Equation S9 into S6 and solving the remaining integral yields the solution for the stationary distribution of *N* in the form of a Gamma distribution with shape parameter *η* and rate parameter *ϰ*:

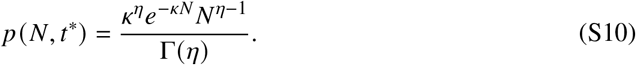

This distribution exists, and thus the stationary distribution of population size can be determine, when *η* > 0, which leads to the following condition:

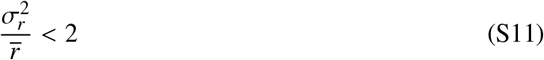

If this condition is not satisfied, the distribution collapses to a Dirac-Delta function centered on *N* = 0. This implies that population extinction occurs rapidly and therefore we denote inequality S11 as the **strong limit** for persistence. Within the strong limit, *p* (*N, t*^*^) is a monotonically decreasing function of *N* when 0 < *η* < 1 with the mode of the distribution 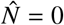. When the *η ≥* 1, *p* (*N, t*^*^) is a unimodally distributed along *N* with the mode given by:

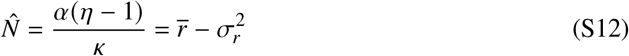

The distinction between the behavior of *p* (*N, t*^*^) around *η* = 1 has a strong influence on extinction risk because

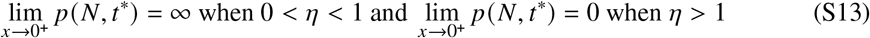

implying that extinction is far less likely an outcome when *η* > 1.

For the classic *r − K* logistic model with variation in *r* only, May (*10*) posited that extinction would be a likely outcome whenever the mean population size was less than half its deterministic carrying capacity, *K*. This corresponds to the point where the probability of distribution of *N* begins to accumulate around zero and *K* (note that *K* is an absorbing state in this model, whereas the deterministic carrying capacity, 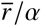, is not absorbing in the *r − α* model). The mean population size for the *r − α* model is

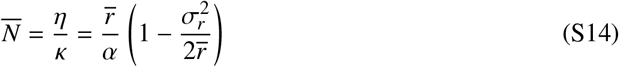

and mean population size is reduced to less than half of the deterministic carrying capacity when

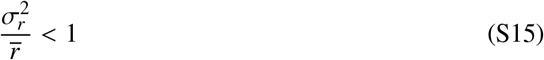

which corresponds precisely to *η* < 1. We call this boundary, where *η* = 1, the **weak limit** for persistence.

## Appendix B Extinction time in the *r − α* logistic model with variation in *r*.

Lande (*12*) calculated the mean time to extinction for a Logistic-type model where the population grew exponentially at rate *r* up to a ceiling given by the carrying capacity. Assuming environmental stochasticity in *r*, he showed that the scaling of extinction time with carrying capacity was logarithmic for 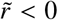, near linear (the square of the logarithm) for 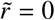, and according to a power-law for 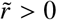, where 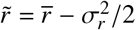 is the long run growth rate of the population.

Using the same approach taken by Lande (*12*) (also described in Otto and Day (*41*)), the formula for the mean waiting time until absorption, given a starting density *N*_0_, when the diffusion process has only one absorbing (extinction) state is:

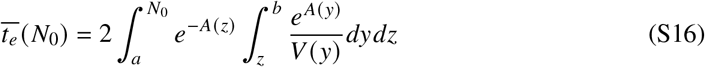

where 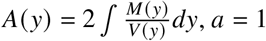 is the extinction boundary, and *b* = ∞ is an upper boundary value for the population size. For the *r − α* model, the functions *M* (*y*) and *V* (*y*) given in Equation S8 do not permit a particularly useful analytical solution of the integral except at the weak limit for persistence where 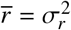, in which case:

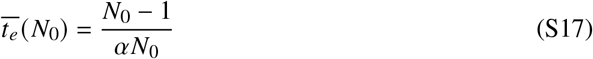

Here, for a reasonably large *N*_0_, the mean time to extinction is well approximated as 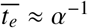. This means that the weak persistence boundary represents a division of populations whose mean time to extinction either exceeds (for 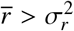) or is less than (for 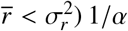. Numerical integration of Equation S16 shows that even small deviations away from the weak persistence boundary generate rapid changes in the mean time to extinction (figure S1). Furthermore, we show that the mean time to extinction declines with *α* according to a power-law, with slope determined by the ratio 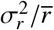 (figure S2). This is consistent with Lande’s (*12*) observations.

The time units of 1/*α* are most easily envisioned in units of generations; given that the mean generation time of a population is 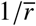, the mean number of generations until extinction at the weak persistence boundary is given by 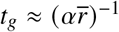. Thus, along the weak persistence limit, populations with the faster rates of intrinsic growth will traverse a larger number of generations before extinction but the absolute time to extinction will not vary.

## Appendix C Simulating the stochastic *r − α* model

We performed stochastic simulations of the *r −α* model using the method proposed by Milstein (*37*). Using this method, Equation S4 can be simulated as:

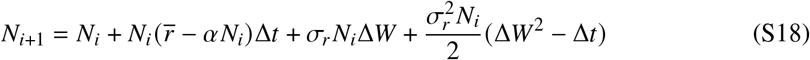

where Δ*t* is the length of the time step, *i* is an index variable, and Δ*W* is an IID random variable drawn from a normal distribution with zero mean and standard deviation equal to 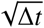. The Milstein method offers superior strong-order convergence over the Euler-Maruyama method through the addition of the last term in equation S18. For all simulations, we set Δ*t* = 0.01. We set the initial population size for simulations as 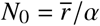. We contrasted this choice with simulations where the initial size was drawn randomly from *p* (*N, t*^*^) (the Gamma distribution given by equation S10), but found no systematic differences. We assumed extinction to have occurred if *N* < 1. Simulations were performed in *Mathematica* version 13.3.1. Although *Mathematica* does have built in functionality for simulating stochastic differential equations, we preferred to directly simulate equation S18 so that a stop condition (extinction) could be embedded for improved efficiency.

## Appendix D Model Simulations

### Simulating the stochastic temperature-dependent *r* (*T*)*−α* model

We constructed a temperature-dependent version of the *r − α* model using the framework described in Vasseur (*8*). Here, the simplest and most generic assumption is that the strength of density-dependence (*α*) does not vary as a function of temperature. This corresponds to a scenario in the *r − K* Logistic framework where both *r* and *K* vary in a perfectly correlated fashion (*8*). Recent work has explored variations on this assumption (*24, 42*) and experimental work using *Daphnia* suggests a doubling of the strength of density dependence for every 7°C of warming (*29*). However, to maintain tractability in our model we assume that *α* is independent of temperature.

Although many mathematical forms of the TPC have been proposed, we utilize the form of the TPC that matches our experimental population of *Paramecium caudatum* (see below) in all of our simulation models. This TPC, fit using the “lactin2” model, is described in detail in Appendix E. Notably, this form of the TPC has the classic left-skewed unimodal shape and it produces negative values at both temperature extremes, consistent with what is known about the thermal performance of the intrinsic growth rate *r*. We also tested other forms of the TPC in our models, including that used in (*42*) and found results consistent with those presented here.

Turning now to the challenge of incorporating variation in temperature into the *r − α* model, it is clear that we cannot write an auxiliary process of the form 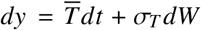 that would translate into an Itô-diffusion process due to the exponential functions in the thermal performance curve, *r* (*T*). Seemingly, any reasonable mathematical representation of the thermal performance curve makes it difficult, if not inadmissible, to being written as a simple diffusion process due to the way that non-linear terms are typically combined in TPCs. Moreover, it is certain that such a formulation would have no analytical utility beyond providing a framework for numerical simulation of the model. Instead, we numerically simulate the temperature dependent *r − α* model using a hybrid approach where the population and temperature dynamics are written as an ordinary and stochastic differential equation pair (shown in Equation S1).

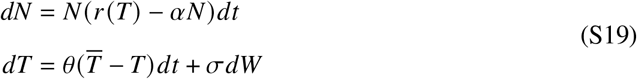

Here, temperature dynamics are generated using an Ornstein-Uhlenbeck (OU) process with mean temperature 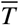, reversion speed *θ*, and volatility *σ*. Given that the variance of the OU process depends upon *σ* and *θ*, we set 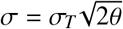, where *σ*_*T*_ is the desired standard deviation of the resultant distribution of temperature. Although our work incorporating variation directly in *r* assumed no autocorrelation in temporally successive values, the downside of being unable to formulate this model as an Itô process means that we cannot avoid the introduction of some autocorrelation into successive values of *r* (*T*). However, we can critically evaluate our choice of *θ* so that any effect of temporal autocorrelation on population dynamics should be minimal.

Previous work has shown that increasing autocorrelation leads to a tighter relationship between population size *N*_*t*_ and the equilibrium population size at the current environmental state, here *N*^*^ = *r* (*T*)/*α*. This is known as ‘tracking’ and can be measured by the correlation of the two variables. In circumstances where the equilibrium size is zero (or near zero) within the range of environmental states that are encountered, autocorrelation increases the likelihood of extinction through this tighter relationship. Research using population dynamic models has shown that environmental variation operating at frequencies below *λ*, the eigenvalue of *∂* (*dN*/*dt*)/*∂N*, generates increasingly stronger tracking while frequencies above *λ* do not. For the *r − α* model, the eigenvalue is *r* and thus the potential to track the environmentally determined equilibrium changes as a function of temperature leading to potentially important asymmetries among cold and warm periods (see (*24*)).

The autocorrelation function of the OU model decays according to *e*^*−θk*^, where *k* is the time-lag. We set *θ* = 5, which produces a temporal autocorrelation that decays rapidly over short time-scales and, importantly, to an insignificant level within *k* = 1 time unit – which corresponds roughly to the generation time of the population (1/*r*) near the optimum value of *r* (*T*). As the magnitude of *r* (*T*) declines toward the upper and lower critical temperatures *T*_min_ and *T*_max_, the impact of autocorrelation weakens as the critical frequency for tracking drops, making a larger fraction of the variation less consequential. Any autocorrelation introduced through use of the OU process should therefore have little impact on population dynamics and extinction. We leave a systematic investigation of the role of autocorrelation in temperature for future work.

Assuming that temperature is given by an IID normally distributed random variable 𝒯 = 𝒩 (*μ*_*T*_, *σ*_*T*_), then *r* (𝒯) is also an IID variable from the transformed distribution of 𝒯. If the function *r* (*T*) is linear, then the mean and variance of the transformed distribution can be easily calculated knowing *r* (*T*) and *r*^′^(*T*). However, if the function is non-linear, the mean and variance of *r* (𝒯) will depend on both *μ*_*T*_ and *σ*_*T*_ (due to Jensen’s inequality) and all of the non-zero higher derivatives of *r* (*T*). In addition, higher-order moments (e.g. skewness and kurtosis) will be introduced into the transformed distribution of *r* (𝒯), leading to radically different shapes and properties (see figure). Given these differences in the distribution of *r* (𝒯) relative to those for which the analytical limits for persistence were established in Appendix A, we can ask how well the weak and strong limits for persistence predict outcomes under temperature variation (rather than for direct variation in *r*).

The weak and strong limits for persistence can be determined as functions of the mean and variance of temperature using numerical methods to approximate 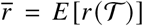 and 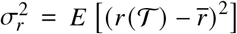. Higher moments of skewness and kurtosis can be similarly determined. We used the *NExpectation* function in *Mathematica* to compute the first four moments across a full range of the independent variables *μ*_*T*_ and *σ*_*T*_. These are shown in figure S3.

We then evaluated the utility of the weak and strong persistence limits by simulating Equation S19 across this same range. We simulated this equation using the Euler-Maruyama method – which is equivalent to Milstein’s method for this model – as:

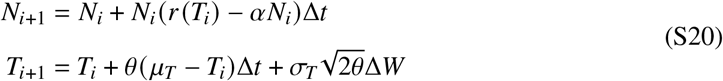

where Δ*W* is a random variable drawn from a normal distribution with mean zero and standard deviation 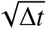. As in simulations of the simpler *r − α* model (see appendix C), we set Δ*t* = 0.01 and we assumed an extinction had occurred if *N*_*i*_ < 1.

### Tailoring a stochastic temperature-dependent *r* (*T*) *− α* model to our experimental system

To more closely replicate our experimental systems, we generated a continuous-time Markov-chain version of our population model which we simulated with the Gillespie stochastic simulation algorithm (*39, 43*). This model/algorithm allowed us to incorporate several processes that are missing in our ODE model but central to our experiments, including demographic stochasticity, temporal autocorrelation of temperatures, and decaying culture conditions.

To accomplish this, we first decomposed the thermal performance curve *r* (*T*) for *P. caudatum* (see below) into a pair of exponential functions representing birth *b*(*T*) and death *d* (*T*) as in (*7, 43*) and then added linear density dependence to the death rate only. This allowed us to reconstruct the *r* (*T*) *− α* model as a birth and death processes. We set the strength of density dependence *α* to generate a maximum population size of 5000 individuals at the beginning of the experiment (consistent with our other models and experimental observations) and assumed a linear increase in *α* over time, resulting in a doubling of *α* by the end of day 56.

Briefly, we simulated this model across a range of mean and variance of temperatures by translating and scaling the 28 day (56 step) time-series that was used in our experiments. Starting from an initial population size of 5000 individuals we randomly determined the next event as a birth or death with weighted probabilities given by *b*(*T*) and *d* (*T*, *N*) and updated the population size accordingly. We advanced time by a random increment drawn from an exponential distribution with mean 1/(*b*(*T*) + *d* (*T*, *N*)) *N*, and we adjusted the temperature accordingly as time advanced past each 12-hour increment. We assumed that an extinction occurred if the population size *N* dropped below 10 individuals, which was the detection threshold used in our experiment. We replicated the result 20 times at each combination of environmental parameters.

## Appendix E Experimental Methods and Results

For our experimental work, we used *Paramecium caudatum* (obtained from Carolina Biological Supply). *P. caudatum* were grown in medium made from spring water (obtained from a natural spring in Roxbury, CT, 41°31’07.7”N, 73°15’45.2”W) autoclaved with 1 g/L of Timothy Hay (Carolina Biological Supply) suspended in a cloth tea bag, which was removed after the water cooled. Bacteria (*E. coli* K12, *Bacillus cereus, Serratia marcesens*) was added to sterile medium 3-4 days prior to the addition of *P. caudatum*. Cells were counted using Leica M165C and M125C stereo dissecting microscopes.

### Measurement of Thermal Performance Curve

We measured the thermal performance curve of *P. caudatum* by growing 12 replicate populations in 6 different constant temperatures (18, 22, 24, 26, 18 and 30°C) in a Percival I-30VL incubator. We initialized each replicate by pipetting between 4 and 8 cells into 1.5 mL of bacterized medium using a 24-well plate. Plates were incubated for between 42 and 68 hours and final cell counts were taken. Population growth rate for each replicate was determined according to:

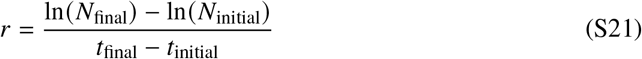

We then fit the “lactin2” model (*44*) to the data in *R* using the *rTPC* package (*38*). The lactin2 model is given by:

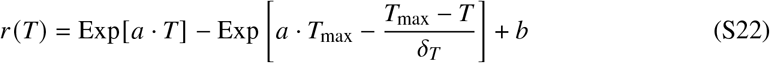

Fitted parameter values statistics are given in Table S1. The fitted curve and data are shown in Figure 4 and is the basis for all our model simulations. Other models fit the data similarly well; however, we chose to use the lactin2 model because it is continuously differentiable, which facilitates taking numerical expectations of the mean and higher moments given a normal distribution of temperature.

### Extinction Experiment

We conducted an experiment to test the utility of the analytical persistence boundaries for predicting extinction in *P. caudatum*. Given the position of these boundaries in the space defined by the mean and variance of temperature, which we established by taking numerical expectations to determine 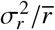, we placed 20 temperature treatments such that treatments spanned the analytical boundaries. Because we had only three incubators that could be programmed to follow a sequence of temperatures (Percival I-30VL outfitted with Intellus Ultra Controllers), we arranged treatments in groups of 5 at low, moderate, and high variance and we used a fixed temperature incubator (Percival I-36LL) for the constant temperature group. Within each incubator, we generated treatments with different mean temperatures by growing cultures in aluminum “chilling” blocks (herein referred to as “blocks”) (Research Products International) that each house 12, 15 mL culture tubes. We attached 2 silicone electric heating pads to each block (12 V, 25 W, and measuring 80 mm x 100 mm) on opposite sides and we wired them to a variable DC power supply, which we used to heat each block a fixed number of degrees above the ambient temperature (see S4). We generated a calibration curve to measure the relationship between voltage and degree of warming and used this to set the voltage outputs needed to achieve each increment used in the experiment. We arranged “chilling blocks” so that within each variance treatment, we had blocks with 0, +3, +6, +9 and +11 °C of warming above the ambient conditions in the incubator. For consistency across different variance treatments, we tied all blocks with the same warming increment to a single power supply.

**Table S1:** Parameter estimates and statistics for lactin2 model TPC (equation S22) fit to *P. caudatum*.

| Parameter | Estimate | Standard Error | T-statistic | 95% CI (low) | 95% CI ( high) | p-value |
| --- | --- | --- | --- | --- | --- | --- |
| $a$ | 0.044 | 0.0373 | 1.18 | -0.0304 | 0.118 | 0.24 |
| $b$ | -1.77 | 0.613 | -2.89 | -2.998 | -0.551 | 0.0051 |
| $T_{\max}$ | 35.3 | 3.34 | 10.56 | 28.594 | 41.913 | 5.53e-16 |
| $\delta_T$ | 5.43 | 6.41 | 0.85 | -7.353 | 18.222 | 0.40 |

In the low, moderate, and high variance treatments, temperatures changed every 12 h, which our previous work had shown was sufficient to generate a response in the population dynamics, but not so long as to overemphasize the impact of singular extreme events. We ran the experiment for 28 days, thus requiring temperature sequences of 56 steps in length. To generate these sequences, we began by creating a normal distribution of 56 temperatures using inverse sampling from the cumulative distribution function of the standard normal distribution at equally spaced intervals {1/57, 2/57, …, 56/57}. This ensured that our small sample of temperatures very closely represented a normal distribution. We then randomly permuted this sequence to form a baseline random sequence that we then scaled and transformed to have (*μ*_*T*_, *σ*_*T*_)= (22,2), (20,3.5), (18,5) for the low, moderate, and high variance treatments respectively. This technique ensured that all treatments experienced the same rank-ordering of temperatures, but had different means and variances. Incubators were set to ‘step’ mode so that temperature changes were discrete; we observed that our blocks took between 60-90 minutes to equilibrate to each step change.

Eight days prior to the start of temperature fluctuations, we inoculated culture tubes with 10 mL of fresh bacterized medium and incubated tubes in blocks at 26°C. Six days prior to the start of the experiment, we added 1 mL from a stock culture of *P. caudatum* (made from the same culture medium) and one sterile wheat seed. The density of cells in the stock culture was 21 cells/mL. Inoculated cultures were incubated at 26°C. Two days prior to the start of the experiment, we counted cells in a 0.25 mL sample from each replicate to ensure that populations were viable. We found a small number of cultures that had zero cells and resampled those on the day temperature fluctuations began. We found three replicates that still had zero cells in a 0.5 mL sample and in these replicates we replaced 5 mL with medium and cells from a stock culture.

Every 7^th^ day of the experiment, we mixed the cultures by gently inverting the tube twice and assayed for presence/absence of cells by microscopy in a 0.25 mL sample. If no cells were present in the initial sample, we took a second 0.25 mL sample to increase our opportunity to find cells. After sampling, we topped each replicate back up to the original 11 mL with sterile medium to compensate for evaporation and losses due to counting. Although we did not directly count cells in dense cultures, we estimate that the maximum abundance in any culture was approximately 100 cells/0.25 mL, leading to a maximum population size of approximately 4400 cells in the 11 mL culture.

We measured the temperatures of each aluminum block daily using an infrared “gun” thermometer (Cole-Palmer). We realized that midway through the experiment, our blocks were not reaching the expected temperatures, potentially due to the heat load placed on our incubators and the compensating effect of increasing cooling in the incubator. We found that on average our warmest blocks (+9 and +11°C) were under-performing by approximately 2.5°C. Rather than alter the design of our experiment midway through, we let the first 28 day cycle complete, and then ran the experiment for a second 28 cycle, repeating the same programmed temperature sequences but with adjusted voltage outputs to reach the intended extent of warming. We censused the experiment only once more at the conclusion of the second 28 day interval.

15 of our experimental replicates did not register a positive density of cells throughout the entire experiment so we discarded those replicates from the analysis. After the first 28 days, 19/225 replicates had gone extinct in our experiment, but with no clear pattern (due likely to the under-performance of our heating apparatus; figure S5). After the second 28 days, 84/225 replicates had gone extinct (65 more extinctions; see Figure 4). We tested for statistically significant differences in the extinction rate across the two groups of treatments separated by the extinction boundary by grouping all of the +9 and +11 °C treatments together and asking if the fraction of observed extinctions was greater than the rate across the whole experiment. We observed 71 extinctions in the 88 non-discarded replicates in this group. We resampled the outcomes of 88 replicates randomly from the 225 non-discarded outcomes 10^7^ times to generate a null distribution for this group. The largest value we obtained from this random sampling was 50, indicating strong significance of our result (*p* < 10^*−*7^).

**Figure S1:**
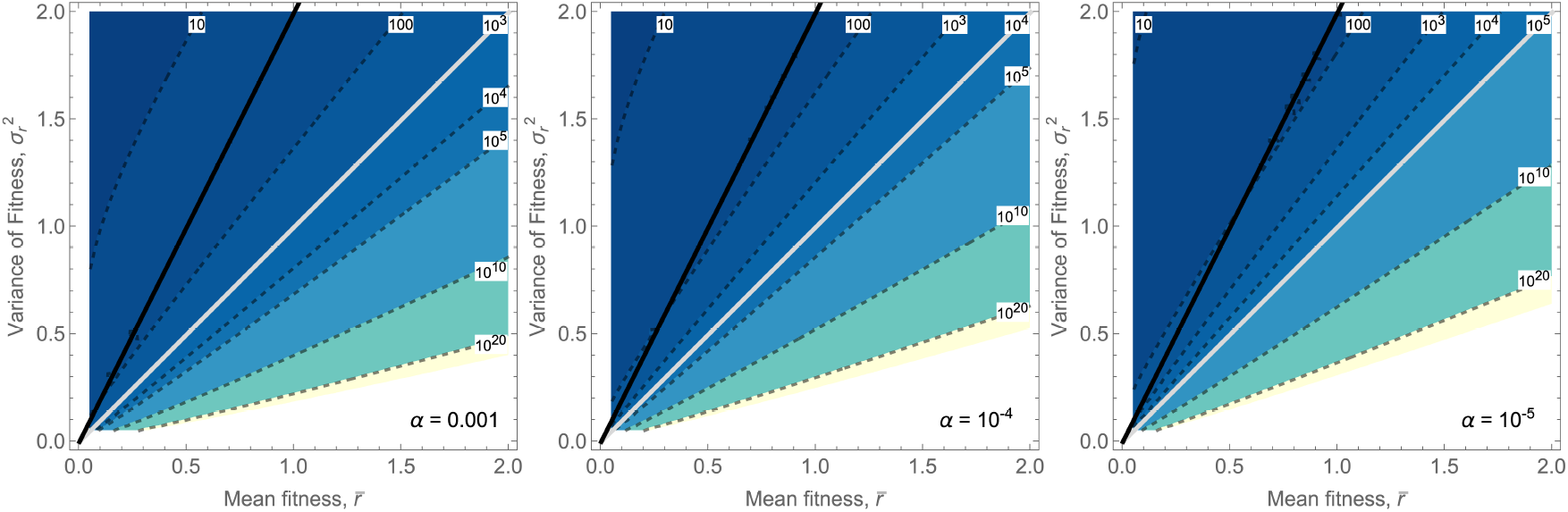
The mean time to extinction for three different values of *α* generated by numerical integration of equation (S16) using an initial population size 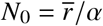. The strong persistence limit 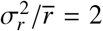 is shown in black and the weak persistence limit 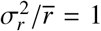 is shown in gray. The weak persistence limit overlies the contour where 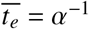.

**Figure S2:**
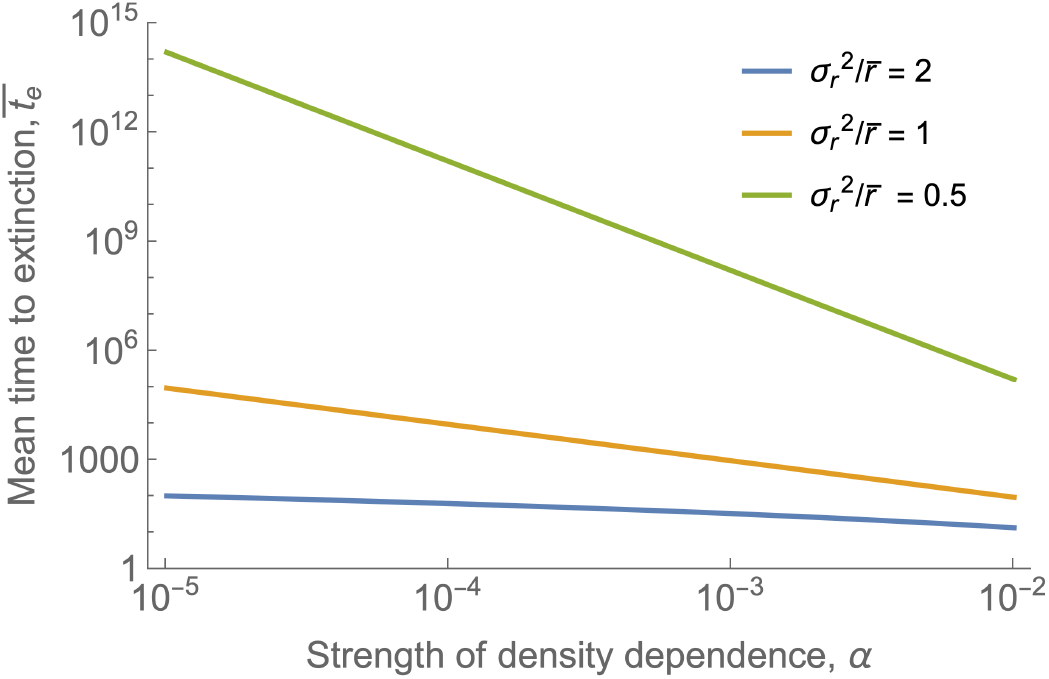
The mean time to extinction as a function of the strength of density dependence *α* for three different values of 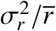.

**Figure S3:**
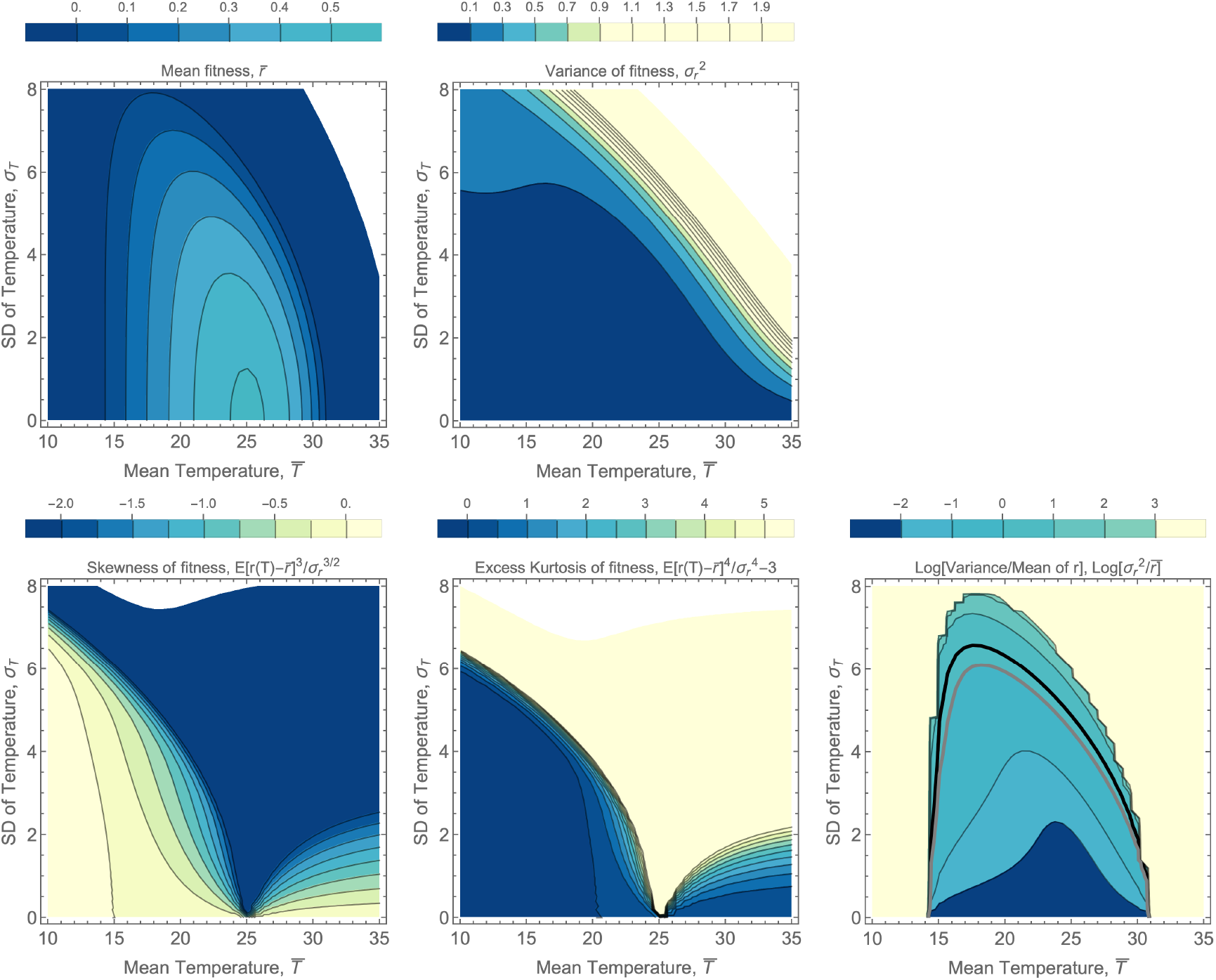
The first four moments of the distribution of r(T) generated by a normal distribution of T with given mean, *μ*_*T*_ and standard deviation, *σ*_*T*_. The final panel shows the ratio of 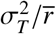 on a logarithmic scale, with the weak and strong boundaries denoted by the gray and black lines. The skewness and excess kurtosis lead to asymmetric and long-tailed distributions over most combinations of mean and SD of temperature. The thermal performance curve used in this figure is that shown in Figure4 and described in Appendix E

**Figure S4:**
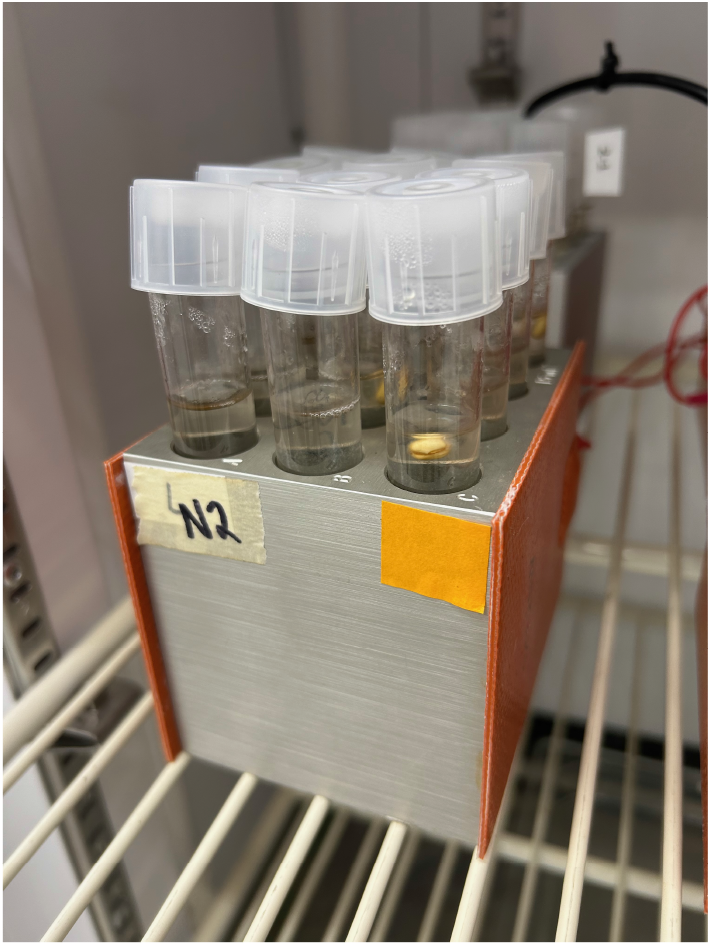
Aluminum ‘chilling’ block with 12, 15 mL culture tubes and 2 surface mounted 12 V (25 W) silicone heating pads. Tube caps are loosely fitted to allow gas exchange.

**Figure S5:**
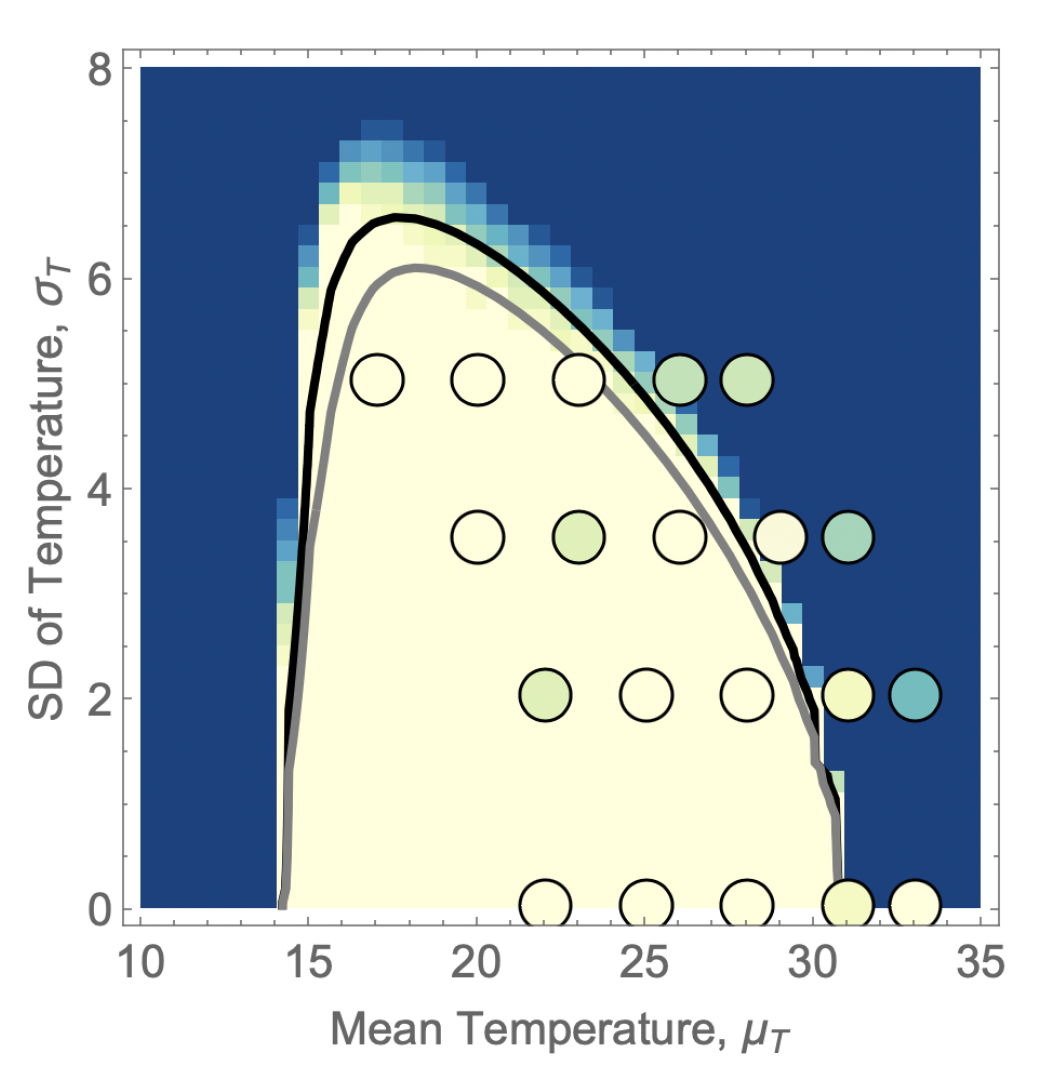
The fraction of experimental replicates extinct after 28 days (shown as bubbles superimposed on the background of our SDE model outcomes). Here our warmest blocks underperformed their target values by approximately 2.5°C; the locations of bubbles in the plot is the “expected” rather than the “observed” locations for the treatments.

**Caption for Movie S1. The fraction of replicate simulations of the stochastic** *r − α* **model persisting through time at each combination of mean and variance of fitness** 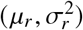. Given the large variation in time-to-extinction across this parameter space, the time step between frames increases by a factor of 10 every *log*_10_ interval. This is an animated version of Figure 1 from the main paper with the same model parameters except that here only 20 replicates per grid cell were simulated.

**Caption for Movie S2. The fraction of replicate simulations of the stochastic** *r* (*T*) *− α* **(temperature dependent) model persisting through time at each combination of mean and standard deviation of temperature** (*μ*_*T*_, *σ*_*T*_). Given the large variation in time-to-extinction across this parameter space, the time step between frames increases by a factor of 10 every *log*_10_ interval. This is an animated version of Figure 3 from the main paper with the same model parameters except that here only 20 replicates per grid cell were simulated.

